# Natural cryptic variation in epigenetic modulation of an embryonic gene regulatory network

**DOI:** 10.1101/831099

**Authors:** Chee Kiang Ewe, Yamila N. Torres Cleuren, Sagen E. Flowers, Geneva Alok, Russell G. Snell, Joel H. Rothman

## Abstract

Gene regulatory networks (GRNs) that direct animal embryogenesis must respond to varying environmental and physiological conditions to ensure robust construction of organ systems. While GRNs are evolutionarily modified by natural genomic variation, the roles of epigenetic processes in shaping plasticity of GRN architecture are not well-understood. The endoderm GRN in *C. elegans* is initiated by the maternally supplied SKN-1/Nrf2 bZIP transcription factor; however, the requirement for SKN-1 in endoderm specification varies widely among distinct *C. elegans* wild isotypes owing to rapid developmental system drift driven by accumulation of cryptic genetic variants. We report here that heritable epigenetic factors that are stimulated by transient developmental diapause also underlie cryptic variation in the requirement for SKN-1 in endoderm development. This epigenetic memory is inherited from the maternal germline, apparently through a nuclear, rather than cytoplasmic, signal, resulting in a parent-of-origin effect (POE), in which the phenotype of the progeny resembles that of the maternal founder. The occurrence and persistence of POE varies between different parental pairs, perduring for at least ten generations in one pair. This long-perduring POE requires piwi-piRNA function and the germline nuclear RNAi pathway, as well as MET-2 and SET-32, which direct histone H3K9 trimethylation and drive heritable epigenetic modification. Such non-genetic cryptic variation may provide a resource of additional phenotypic diversity through which adaptation may facilitate evolutionary changes and shape developmental regulatory systems.

## Introduction

The “Modern Synthesis” of the early 20^th^ century articulated how biological traits shaped by Darwinian forces result from random mutations following the rules of Mendelian inheritance (1). Since that formulation, it has become clear that non-genetic heritable mechanisms can underlie substantial differences in traits between individuals (reviewed in ref. 2). Extensive epigenetic reprogramming occurs in the germline and in gamete pronuclei after fertilization to maintain the totipotent state of the zygote. In mammals, disruption of this process often leads to lethal consequences (3, 4). In *C. elegans*, aberrant reprogramming of epigenetic memory can result in transgenerational accumulation of inappropriate epigenetic marks and a progressive sterile mortal germline (Mrt) phenotype (5, 6). In many cases, the Mrt phenotype is exacerbated by heat stress, demonstrating that environmental factors may influence epigenetic reprogramming in the germline and that these epigenetic modifications may be passed to subsequent generations (7, 8). Interestingly, *C. elegans* wild isotypes, each carrying a unique haplotype, exhibit variation in the temperature-induced Mrt phenotype, suggesting differential stress response and distinct epigenetic landscapes in natural populations of the species (9).

Many of the documented instances of epigenetic inheritance in mammals are parental or *intergenerational* effects (no more than three generations for female transmission and less than two generations for male transmission), which can be attributed to direct exposure of the developing embryos *in utero* to the triggers that alter epigenetic states (2). Parental traumatic experience can trigger heritable behavioral changes, and nutritional status of the parents can cause metabolic remodeling in the offspring, which often lasts for one or two generations (e.g., refs. 10, 11). Epidemiological analyses on different human cohorts demonstrate that paternal grandfather’s food access during pre-puberty period affects the mortality of the grandsons (12–14), revealing the potential for long-term *transgenerational* epigenetic inheritance (TEI) that is induced by environmental conditions.

Studies over the past decade on *C. elegans* have provided convincing evidence for TEI that persists for at least three generations. Small RNAs are prime candidates for mediators of epigenetic memory, as their expression undergoes only minimal reprogramming in the germline and embryos (15, 16). Primary siRNAs, processed from exogenous dsRNAs or endogenous small RNAs by DICER, are loaded onto the RNA-induced silencing complex (RISC) and mediate degradation of mRNA targets, thereby silencing gene expression. In addition, primary siRNAs, including PIWI-interacting RNA (piRNA – 21U) in the germline, can guide RNA-dependent RNA polymerases (RdRPs) to particular target mRNAs and then amplify silencing signals by producing an abundance of secondary 22G siRNAs. In the germline, HRDE-1 binds to these secondary siRNAs and localizes to the nucleus, where it recruits NRDE-1/2/4 to the nascent transcripts and genomic sites targeted by the small RNAs. This complex then inhibits RNA polymerase II elongation and directs deposition of repressive H3K9me3 marks on the corresponding genomic loci, mediated by histone methyltransferases MET-2 (H3K9me1/2), SET-23 (H3K9me1/2/3) and SET-32 (H3K9me3). Amplification of secondary siRNAs by RdRPs prevents loss of epigenetic memory over multiple generations and, therefore, may permit long-term heritable epigenetic responses (reviewed in refs. 17, 18).

We have uncovered natural epigenetic variation in the modulation of the gene regulatory network (GRN) that directs embryonic endoderm development in *C. elegans*. The maternal SKN-1/Nrf2 transcription factor activates the mesendoderm GRN in the EMS blastomere at the four-cell stage. EMS subsequently divides to produce the E founder cell, which gives rise exclusively to the intestine, and its anterior sister, the MS founder cell, which produces much of the mesoderm. A triply redundant (Wnt, MAP kinase, and Src) extracellular signal sent from the neighboring P_2_ blastomere is received by EMS and acts in parallel with SKN-1 to activate the endoderm GRN in the E lineage. In the wild type laboratory N2 strain, elimination of maternal SKN-1 function causes fully penetrant embryonic lethality and a partially penetrant loss of gut: while the majority of the E cells in embryos lacking SKN-1 adopt the mesoectodermal fate of the normal C founder cell, ~30% undergo normal gut differentiation as a result of this parallel signaling input (SI Appendix, Fig. S1) (reviewed in refs. 19, 20).

We recently found that the requirement for the SKN-1 and Wnt inputs into the endoderm GRN, shows widespread natural variation across genetically distinct *C. elegans* wild isotypes. While removal of SKN-1 in some of the isotypes causes loss of intestine in virtually 100% of embryos, in other isotypes a majority of the embryos still differentiate endoderm. Interestingly, low SKN-1 requirement appears to be partially compensated by a higher MOM-2/Wnt requirement. Thus, the early stages in the endoderm GRN has undergone rapid change during relatively short evolutionary periods in *C. elegans* (21, 22). In this study, we focus on variation in a single major input node, SKN-1.

We report here that, although much of the variation in SKN-1 requirement results from genetic differences between the wild isotypes (21), it is also determined in part by cryptic, stably heritable epigenetic variation. This effect is uncovered from reciprocal crosses between wild isotypes with quantitatively different phenotypes. This parent-of-origin effect (POE) is transmitted exclusively through the maternal germline. When mothers experience dauer diapause, an alternative developmental stage in *C. elegans* that confers resistance to environmental insults and longevity, the POE appears to be transmitted through the maternal nucleus, rather than cytoplasmic factors, and can persist through many generations. We further show that this stress-induced POE requires factors that direct H3K9 methylation and the nuclear RNAi machinery. These findings reveal that heritable epigenetic states can underlie differences between natural wild isotypes and can influence developmental plasticity in early embryos. Such cryptic epigenetic variation provides a potential resource upon which natural selection might act, thus contributing to evolution of GRN architecture (23).

## Results

### Transgenerational parent-of-origin effect alters the SKN-1 dependence of endoderm formation

The requirement for SKN-1 in endoderm specification varies dramatically across *C. elegans* isotypes (21). Depending on the isotype tested, between 0.9% and 60% of arrested *skn-1(RNAi)* embryos undergo gut differentiation when maternal SKN-1 function is eliminated. The behavior of each isotype is quantitatively highly reproducible, showing low variability through many generations, when analyzed by different laboratories and researchers, and from independent lines established from different founder animals (21).

While performing crosses between isotypes with quantitatively different SKN-1 requirements, we found that the outcomes differed depending on the sex of the parent in reciprocal crosses. We initially tested two isotypes in which we observed dramatically different *skn-1(RNAi)* phenotypes: the laboratory N2 strain (29 ± 0.4% sd with gut; n = 1320) and the wild isotype JU1491 (1.2 ± 0.4% sd; n = 1228) (SI Appendix, Fig. S2A, p < 0.001). These results agree well with our previous findings (21). Consistent with variation at the level of maternal components, we found that in reciprocal crosses (i.e., male N2 × JU1491 hermaphrodite, and *vice-versa*), the quantitative requirement for SKN-1 reliably followed that of the maternal line (Fig. 1B). Unexpectedly, however, we found that this non-reciprocality persisted in subsequent generations: the average phenotype of F2 and F3 embryos continued to follow more closely the behavior of their grandmothers and great-grandmothers than their paternal ancestors (Fig. 1C), despite the fact that, with the exception of the mitochondrial genome, the F1 progeny genotypes should be identical regardless of the sex of the founder P0. Thus, these two strains showed a strong parent-of-origin effect (POE) that persists through multiple generations.

**Fig. 1:**
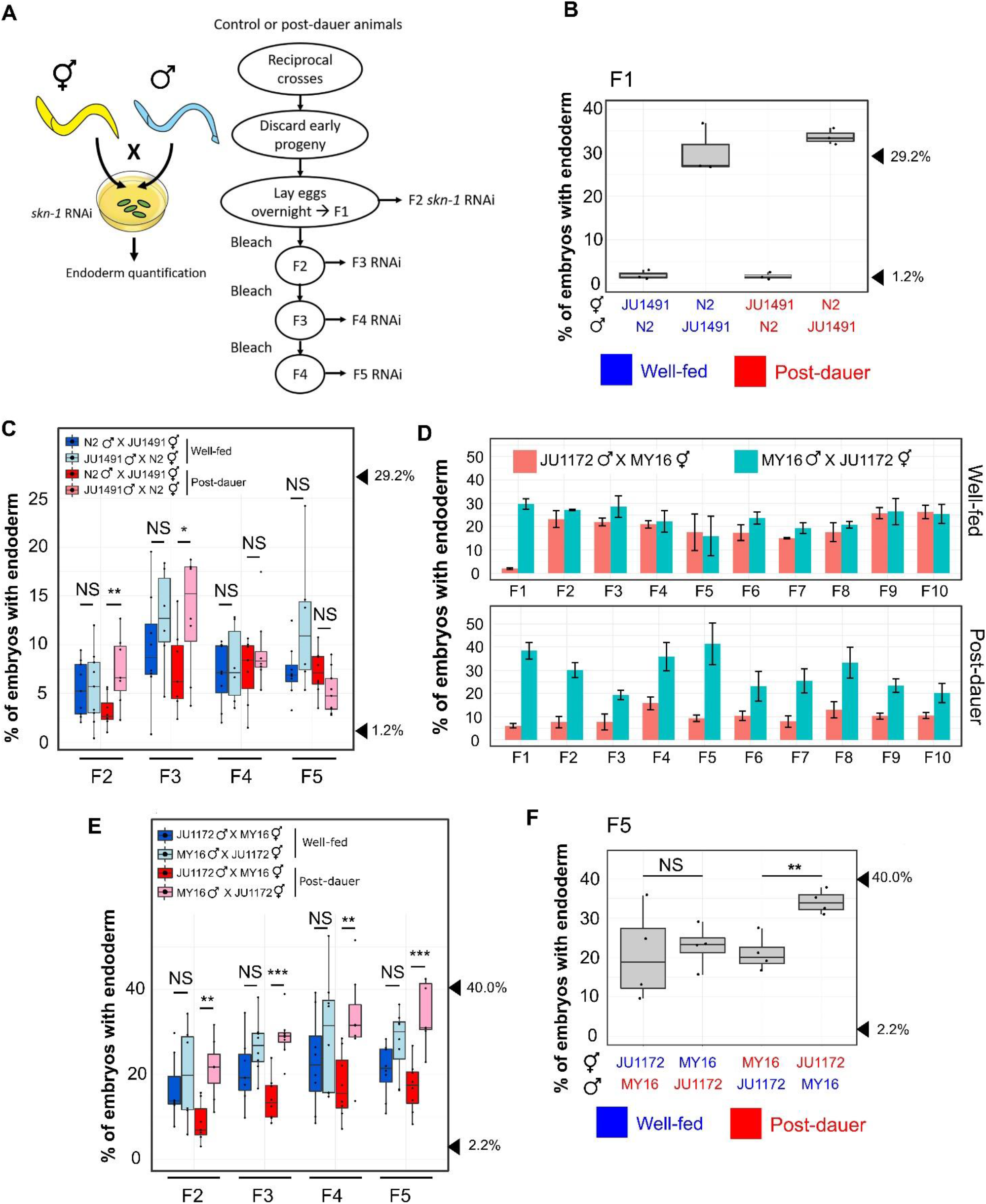
Dauer diapause-stimulated Parent-of-Origin Effect (POE). A) Schematic of the POE assay strategy. Left: to test for a maternal effect, reciprocal crosses using well-fed or post-dauer animals were performed on *skn-1* RNAi plates and the fraction of arrested F1 embryos with differentiated gut was examined. Blue and yellow colors represent different wild isotypes. Right: to study the transgenerational phenotypes, reciprocal crosses were performed using different wild isotypes on NGM plates. Embryos were collected from mated hermaphrodites to establish the F1 population. Day 1 (Fig. 1D) or Day 2 (Fig. 1C, E and F; Fig. 2C; Fig. 4B; SI Appendix, Fig. S4) gravid worms were treated with alkaline hypochlorite solution to obtain the next generation embryos. L4 hermaphrodites were isolated for *skn-1* RNAi assays at each generation (see Materials and Methods). B) Strong maternal effect in F1 embryos from mated *skn-1(RNAi)* mothers in N2 × JU1491 crosses. At least three independent crosses were performed. Arrowheads indicate phenotypes of N2 (29.2%) and JU1491 (1.2%). The phenotypes of the F1 progeny are not significantly different from those of their respective mother in all crosses, regardless of feeding status. One-sample t-test (p > 0.05). C) POE in *skn-1(RNAi)* embryos from four generations: F2, F3, F4 and F5 derived from reciprocal N2 × JU1491 crosses. At least five independent crosses were performed for each treatment. Arrowheads indicate the phenotypes of N2 (29.2%) and JU1491 (1.2%). D) Dauer diapause-induced POE persists for at least 10 generations in JU1172 × MY16 descendants. Three independent crosses were performed. Alkaline hypochlorite treatment was performed on Day 1 gravid adults at each generation. Error bars represent +/− standard error of the mean. Two-way ANOVA (p = 0.08 and 9.14 × 10^−11^ for the well-fed and post-dauer experiments (F2-F10), respectively). E) Robust POE in *skn-1(RNAi)* embryos over four generations: F2, F3, F4 and F5 derived from reciprocal JU1172 × MY16 crosses. At least five independent crosses were performed for each treatment. Alkaline hypochlorite treatment was performed on Day 2 gravid adults at each generation. Arrowheads indicate the phenotypes of JU1172 (40.0%) and MY16 (2.2%). F) POE seen in *skn-1(RNAi)* F5 embryos from reciprocal crosses between MY16 and JU1172. Data points represent independent crosses. Arrowheads indicate the phenotypes of JU1172 (40.0%) and MY16 (2.2%). For panels B and F, blue text represents experiments with well-fed animals, while red text is for experiments with post-dauer animals. Two sample t-test (NS p > 0.05, * p ≤ 0.05, ** p ≤ 0.01, *** p ≤ 0.001). Boxplot represents median with range bars showing upper and lower quartiles.

### Dauer diapause stimulates long-term transgenerational POE through the maternal line

As epigenetic inheritance can be environmentally triggered, it was conceivable that the POE we observed might be influenced by the experience of the parents. Indeed, we found that POE was seen only when the P0 parents had been starved and experienced an extended period (~2 weeks) of dauer diapause. In contrast, the progeny of P0’s that were continuously well fed showed an intermediate average phenotype that was not significantly different between descendants of reciprocal crosses (Fig. 1C), consistent with the known multigenic characteristic of the phenotype (21).

We sought to determine whether this environmentally triggered POE extends to other wild isotypes. Isotypes MY16 (in which only 2.2 ±1% sd (n = 1169) of *skn-1(−)* embryos make endoderm) and JU1172 (40 ± 3% sd; n = 1491) (SI Appendix, Fig. S2B, p < 0.001) show widely different quantitative phenotypes (21). Consistent with our previous findings, in reciprocal crosses of MY16 and JU1172, we observed a strong maternal effect in the requirement for SKN-1: F1 embryos from mated *skn-1(RNAi)* mothers followed the maternal phenotype (Fig. 1D). Once again, in control experiments with well-fed founder P0 worms, this maternal effect quickly dissipated and was not detectable in F2 embryos (Fig. 1D and E). However, when the parental worms experienced dauer diapause, the average *skn-1(RNAi)* phenotype of their descendants reliably followed that characteristic of the maternal line through at least ten generations (Fig. 1D and E – the experiments shown in the two panels were performed using parents of different ages; see below), a strongly perduring effect.

While dauer development enhances POE, we found that it is not an absolute requirement in all cases. Specifically crosses of JU1491 and JU1172 revealed weak POE even without the diapause trigger, although the effect was enhanced by dauer development (Fig. 2C). This observation suggests that cryptic epigenetic differences between some natural isotypes may exist even in the absence of an environmental or physiological trigger. Finally, we found that this effect does not appear to be general to all isotype pairs that show very different phenotypes: for example, diapause-induced POE was not detectable with N2 and MY16 (SI Appendix, Fig. S3).

**Fig. 2:**
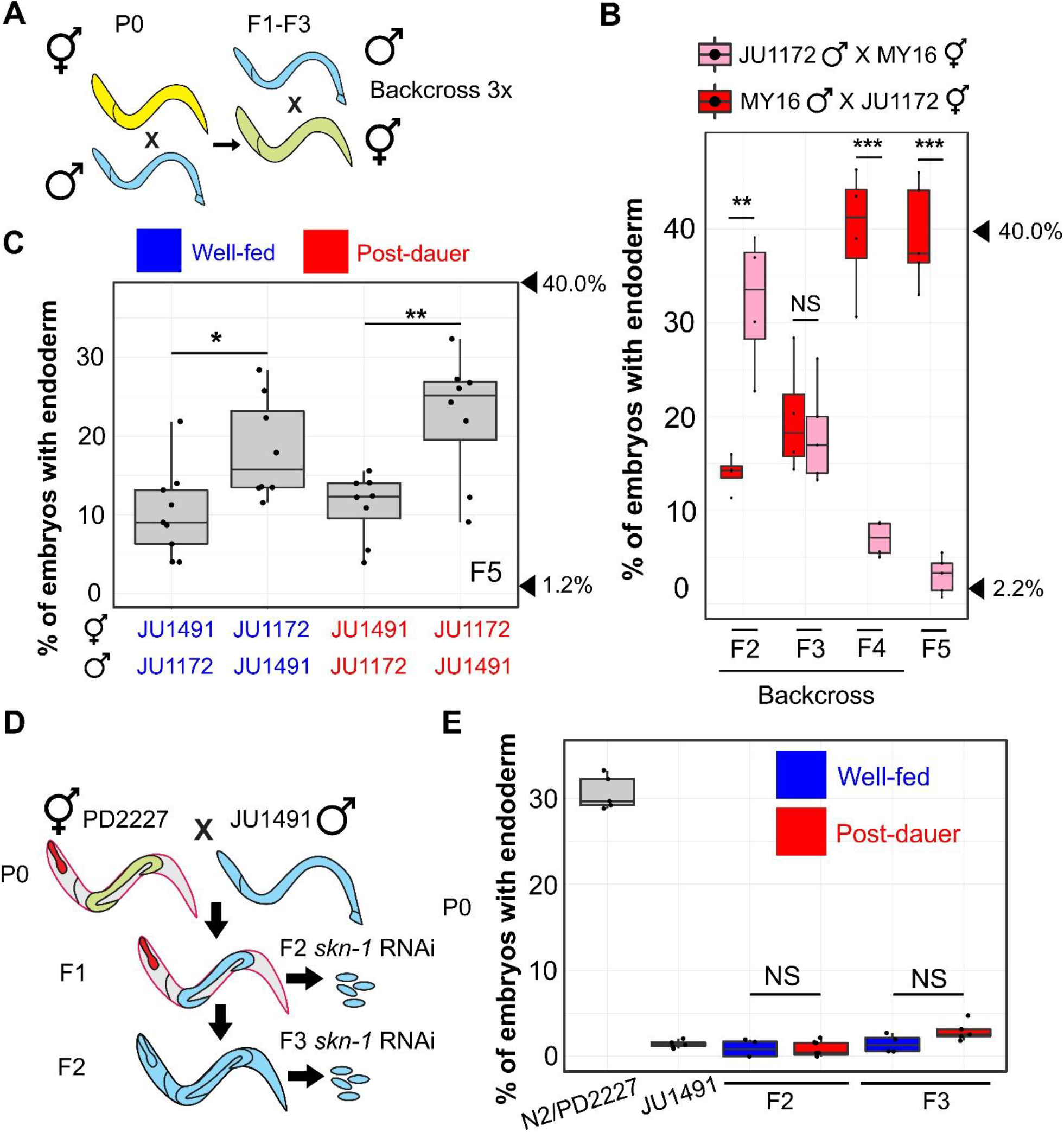
POE is not attributable to mitochondrial DNA or cytoplasmic inheritance. A) Schematic of mitochondrial transfer experiment. Blue and yellow represent different wild isotypes. Five to ten hermaphrodites were used to backcross to the paternal strain at every generation for three generations. B) POE in *skn-1(RNAi)* embryos. Data points represent replicates from a single reciprocal cross using post-dauer animals. Arrowheads indicate phenotypes of JU1172 (40.0%) and MY16 (2.2%). C) POE shown by *skn-1(RNAi)* F5 embryos of reciprocal crosses performed between JU1491 and JU1172. Data points represent independent crosses. Arrowheads indicate phenotypes of original JU1172 (40.0%) and JU1491(1.2%) strains. D) When PD2227 (N2^GPR-1 OE^) hermaphrodites are mated to JU1491, non-Mendelian segregation of maternal and paternal chromosomes result in some F1 mosaic animals that express the maternally derived pharyngeal marker (*myo-2*∷mCherry) but lose the body wall muscle (*myo-3*∷mCherry) and germline (*mex-5*∷GFP) markers. F2 self-progeny of the F1 mosaic animals contain only the JU1491 nuclear genome (see SI Appendix, Fig. S6). E) *skn-1(RNAi)* phenotype of N2/PD2227, JU1491, and F2 and F3 embryos from the mosaic animals. For panels C and E, blue represents experiments performed using well-fed animals while red represents experiments performed using post-dauer animals. Two sample t-test (NS p > 0.05, * p ≤ 0.05, ** p ≤ 0.01, *** p ≤ 0.001). Boxplot represents median with range bars showing upper and lower quartiles.

As expected for successful crosses, in all cases ~50% of the F1 offspring were males (SI Appendix, Fig. S4A and D; one-sample t-test, p > 0.05). Further, cultures established from at least eight randomly selected F1s of successful crosses (with ~50% F1 males) all showed POE (SI Appendix, Fig. S4C), thus ruling out the possibility that the maternal-line bias of the phenotype might result from frequent selfing.

To distinguish between the paternal and maternal contributions to the POE, we starved either the P0 male or hermaphrodite and traced the POE for five generations following reciprocal crosses. These experiments demonstrated that the diapause-induced POE is inherited exclusively through the maternal germline (Fig. 1F; % of F1 males − SI Appendix, Fig. S4B). This stable non-reciprocality cannot be explained by long-perduring maternal factors in the cytoplasm: each animal produces ~250 progeny and after five generations, this would result in a dilution factor of ~10^11^.

It is noteworthy that *skn-1(RNAi)* is highly penetrant in the isotypes used in this study, with 100% *skn-1(RNAi)*-induced lethality observed across all experiments. This suggests that POE reflects *bona fide* variation in the endoderm developmental input (21), although we cannot rule out that RNAi sensitivity might play a minor role.

### Heritability of POE is associated with the maternal nucleus, not heritable mitochondrial or cytoplasmic factors

Dauer larvae and post-dauer adults exhibit a metabolic shift which may reflect changes in mitochondrial function. Further, starvation has been shown to impact mitochondrial structure and function (reviewed in refs. 24, 25). Thus, the observed maternally directed POE results might arise from differences between the mitochondrial genome sequences in the two strains or might be driven by other cytoplasmically inherited factors. Indeed, wild isotypes MY16 and JU1172 contain 13 polymorphisms in mitochondrial protein coding genes (SI Appendix, Table S1), which could alter energy metabolism and stress responses. To test whether the POE we observed is attributable to maternal inheritance of mitochondria with particular genomic characteristics, we performed reciprocal crosses in which progeny were repeatedly backcrossed to the paternal strain to obtain lines with primarily the MY16 nuclear genome and mitochondria from the JU1172 line and *vice-versa* (Fig. 2A). While a strong POE was initially observed in F2 *skn-1(RNAi)* embryos, this effect was rapidly eliminated as more paternal nuclear DNA was introduced. By the F5 generation, the phenotype was indistinguishable from that of the respective paternal strain (Fig. 2B), suggesting that POE is attributable to the nuclear, rather than mitochondrial, genome. Moreover, the transgenerational POE observed with JU1491 and JU1172 (Fig. 2C) is unlikely to be the result of variation in mitochondrial DNA, as these two strains carry identical mitotypes. Collectively, our results suggest that the POE we observe is not likely to be caused by mitochondrial inheritance.

To further assess whether nuclear or cytoplasmic/mitochondrial factors underlie the observed POE, we took advantage of a genetic system that generates germlines containing nuclei derived fully from the paternal line with maternally derived cytoplasm (including mitochondria). In zygotes overexpressing GPR-1 (N2^GPR-1 OE^) (26, 27), which is required to modulate microtubule-based pulling forces, excessive pulling forces cause the maternal and paternal pronuclei to be drawn to opposite poles before nuclear envelope breakdown. This effect generates mosaic embryos in which the anterior daughter (AB) inherits only the maternal chromosomes, while the posterior (P_1_) receives only the paternal chromosomes. These non-Mendelian events can be scored with the appropriate fluorescent markers (Fig. 2D; SI Appendix, Fig. S5) (27). We found that 72% (n = 230) of the viable F1 progeny from crosses of N2^GPR-1 OE^ hermaphrodites with JU1491 males contained an exclusively paternally derived P_1_ lineage. If cytoplasmic maternal factors were responsible for the observed diapause-induced POE, the effect would be expected to follow the cytoplasm of the founder P0 worms in the F1 hybrids. In contrast, however, we found that the *skn-1(RNAi)* phenotypes of F2 and F3 descendants of F1 mosaic animals (those with an N2-derived AB and JU1491-derived P_1_; SI Appendix, Fig. S5; see Materials and Methods) were indistinguishable from that of the JU1491 strain, regardless of the feeding status of the parents (Fig. 2E). This finding suggests that the diapause-induced POE is associated with heritable changes in the maternal haploid nucleus, not heritable cytoplasmic maternal factors, including the mitochondrial genome.

### POE is not the result of competition in fitness or maternal incompatibility

Parental age has been shown to affect progeny phenotypes in *C. elegans* and other organisms (e.g., refs. 28, 29). To test the possibility that the POE is influenced by differences in maternal age, we collected embryos from day-one adults (Fig. 1D) and day-two adults (Fig. 1C, E and F; Fig. 2C). We detected a strong POE in all cases. Moreover, despite large variation in the *skn-1(RNAi)* phenotype that arises from genetic variation, we observed POE in F5 populations that were established from very late broods (the last few progeny) produced by senescent F1 animals (Fig. 3A). These findings indicate that parental age does not contribute substantially to the POE observed.

**Fig. 3:**
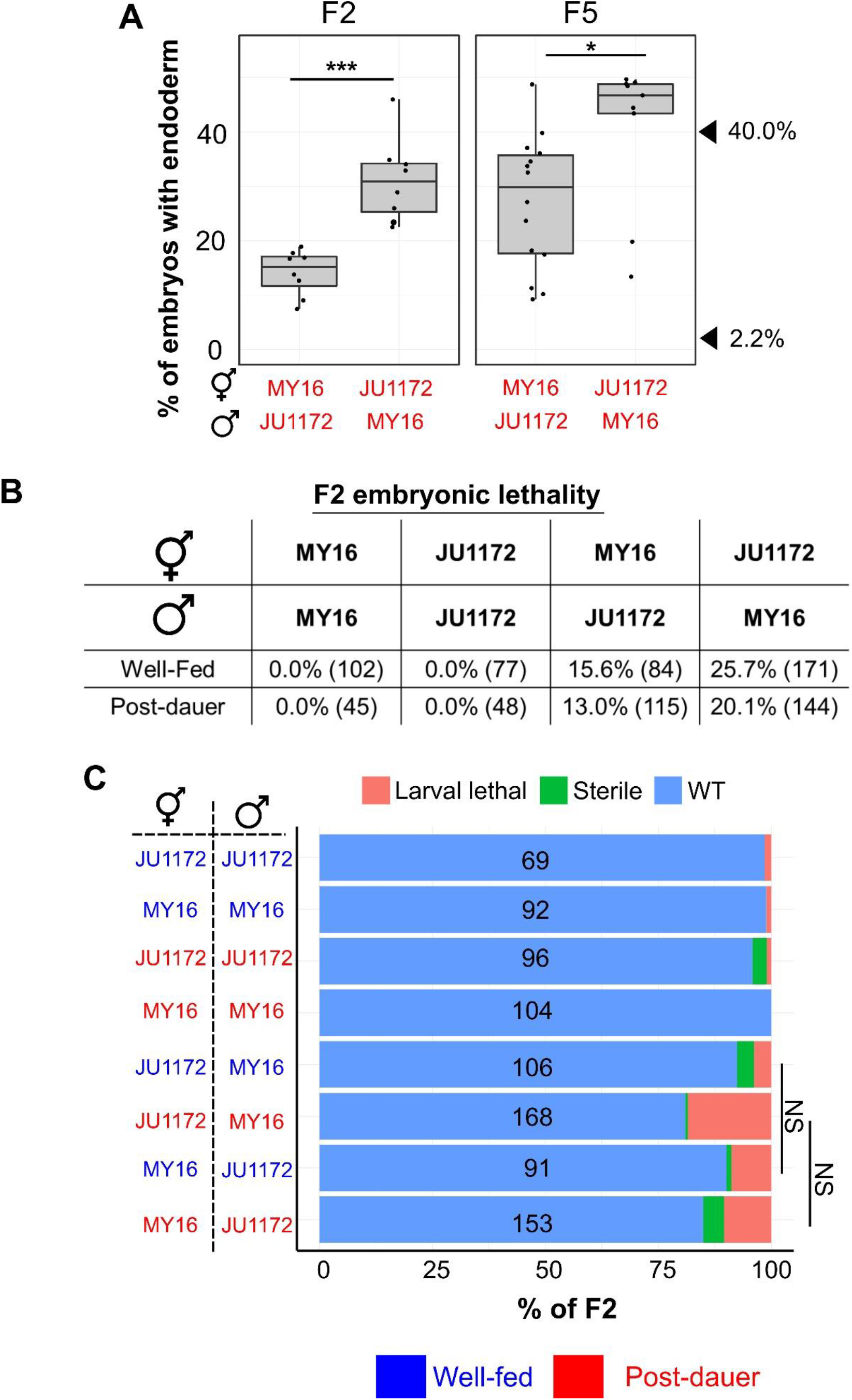
POE is not attributable to competitive fitness or maternal incompatibility. A) Left: POE shown by *skn-1(RNAi)* F2 embryos of reciprocal crosses performed between MY16 and JU1172. Right: Individual data points represent lines derived from late F2s of senescent F1 animals. Two independent crosses were performed for each direction. Two sample t-test (NS p > 0.05, * p ≤ 0.05, ** p ≤ 0.01, *** p ≤ 0.001). Boxplot represents median with range bars showing upper and lower quartiles. The arrowheads indicate the phenotypes of JU1172 (40.0%) and MY16 (2.2%). B) Embryonic lethality in F2s. Two independent broods were examined. Total number of embryos scored is shown in brackets. C) Sterility and larval lethality of F2 progeny. Identity and the sex of the parents are indicated on the left. Total number of animals scored is indicated. Fisher’s exact test (NS p > 0.05). For panels A and C, blue text represents experiments performed using well-fed animal while red text represents experiments performed using post-dauer animals.

The differences in SKN-1 requirement seen as the result of the POE might reflect maternal incompatibility, which favors particular genetic regions as a result of lethality or slow growth (30, 31). If such regions included those known to influence the requirement for SKN-1 in the endoderm GRN (21), there could be selection for the trait following recombination of the two parental genomes. We note that such a possibility would also require that any such selection is triggered only after starvation and dauer development for the cases in which we observed such an essential requirement. Further, we observed strong diapause-induced POE in embryos from *skn-1* RNAi-treated F1 heterozygous mothers, whose genotypes would be identical in the two reciprocal crosses. Thus, the effect at this stage is not attributable to maternal incompatibility resulting in selection against particular allelic combinations that arise by recombination (Fig. 1C-E).

To further investigate whether POE might be driven by genetic incompatibility that is environmentally triggered by starvation/dauer development, we also characterized lethality and fecundity of F2 progeny from the reciprocal crosses. Two mechanisms involving selfish genetic elements that result in maternal incompatibilities were previously described in *C. elegans*: the *peel-1/zeel-1* (30) and *sup-35/pha-1* (31) toxin/antidote systems. The wild isotype JU1172 does not carry the paternal selfish *peel-1/zeel-1* element (30). When mated, MY16 sperm deliver PEEL-1 toxin, causing F2 embryos that are homozygous for the JU1172 *zeel-1* haplotype (~25%) to arrest. We found that, indeed, crosses between JU1172 and MY16 are associated with embryonic lethality. However, although this lethality was slightly lower (~13-16%) when JU1172 was the paternal strain compared to the reciprocal crosses (20-26%; Fig. 3B), the difference is insufficient to explain the strong POE we have observed. Furthermore, this effect does not change appreciably regardless of the experience of the P0 (fed or starved/dauer), which is not consistent with selection induced by this experience. As both MY16 and JU1172 harbor the active *sup-35/pha-1* maternal toxin/antidote element (31), the lethality may reflect an unidentified maternal-effect toxin in the MY16 strain. In addition, the progeny of crosses between MY16 and JU1172 in either direction both showed somewhat reduced fecundity/viability, presumably owing to genomic incompatibility between the two strains (Fig. 3C). However, the parental origin of the P0s did not influence the degree of larval lethality or sterility in the F2 animals (Fig. 3C, Fisher’s exact test p > 0.05). Together these results indicate that genetic incompatibility alone cannot account for the strong POE we observed. Rather, POE appears to result from perduring epigenetic inheritance reflecting the experience of the original founding parents of the cross.

### Maintenance of POE involves the nuclear RNAi pathway and histone H3K9 trimethylation

The findings noted above suggest that POE is mediated through nuclear signals. The nuclear RNAi pathway has been implicated in a number of examples of TEI (7). To assess whether this pathway might be involved in transmitting the POE we have observed, we analyzed the F4 progeny of reciprocal crosses of post-dauer P0’s in which *nrde-4* was knocked down by RNAi (strategy shown in Fig. 4A). While a strong POE was observed in the control animals containing functional NRDE-4, *nrde-4(RNAi)* abrogated the POE in the F5 embryos (Fig. 4B): i.e., the requirement for SKN-1 was not significantly different in the descendants of reciprocal crosses. It was conceivable that this effect might simply reflect a direct role for NRDE-4 in the requirement for SKN-1 *per se*. However, we found that the *skn-1(RNAi)* phenotypes of the MY16 and JU1172 isotypes previously subjected to *nrde-4* RNAi, were indistinguishable from those subjected to control RNAi (Fisher’s Exact test p > 0.05; Fig. 4C). Thus, these findings implicate NRDE-4, and hence the nuclear RNAi pathway, in the POE process.

**Fig. 4:**
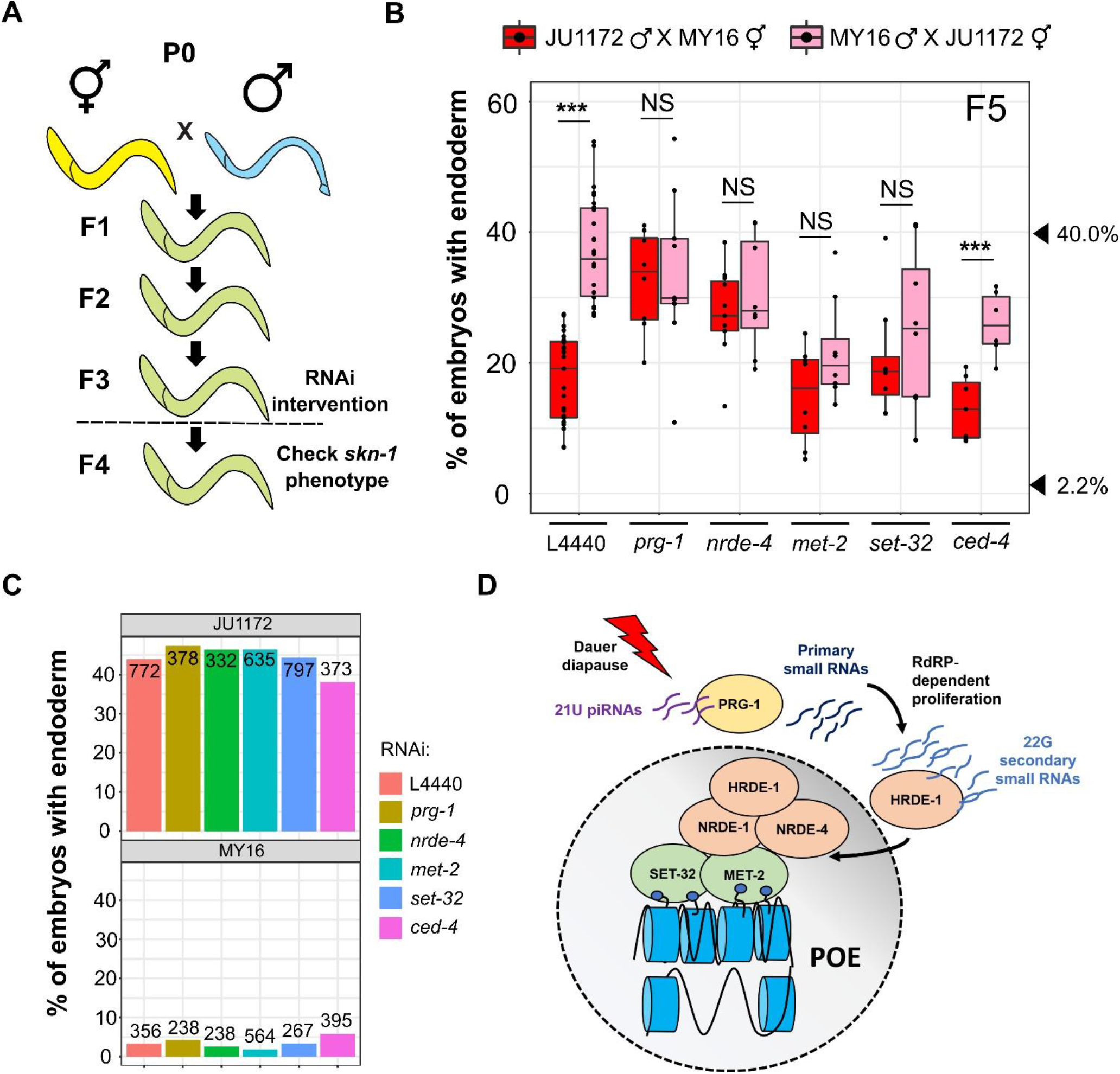
POE requires both the piRNA/nuclear RNAi pathway and factors required for H3K9me3 chromatin marks. A) Schematic of RNAi experiments that test requirement for epigenetic regulators in POE. F3 L4 animals were treated with the indicated RNAis and F4 L4s were used for the *skn-1* RNAi assays. Animals in the control and treatment groups were siblings (see Materials and Methods). B) Knocking down *prg-1*, *nrde-4*, *met-2* and *set-32*, but not *ced-4*, eliminates POE. Data points are replicates from at least two independent crosses. Boxplot represents median with range bars showing upper and lower quartiles. Arrowheads indicate phenotypes of JU1172 (40.0%) and MY16 (2.2%). Two sample t-test (NS p > 0.05, *** p ≤ 0.001). C) The effect of RNAi treatments on the *skn-1(RNAi)* phenotype. MY16 and JU1172 L4s were exposed to L4440 (control), *prg-1*, *nrde-4*, *met-2*, *set-32* or *ced-4* RNAi feeder strains and the *skn-1(RNAi)* phenotype of the F1 were quantified. No difference between different RNAi treatments was detected (Fisher’s exact test, p > 0.05). Total number of embryos scored is indicated. D) Model for POE (see text).

Gene silencing though the nuclear RNAi pathway that results in TEI is mediated through the Piwi-encoding homologue, *prg-1*, and piRNAs that trigger biosynthesis of secondary 22G RNAs (a second homologue, *prg-2*, is likely to be a pseudogene) (32, 33). We knocked down *prg-1/2* in F4 animals from MY16 and JU1172 reciprocal crosses and found that, in contrast to their siblings treated with control RNAi, POE was abrogated in the F5 embryos (Fig. 4B). Control experiments demonstrated that *prg-1/2* RNAi does not affect *skn-1* RNAi efficacy in either parent line (p > 0.05; Fig. 4C). Loss of nuclear RNAi factors lowers the efficacy of RNAi targeting of nuclear-localized RNAs (34–36); however, maternal *skn-1* mRNA in the early embryos is localized in the cytoplasm, and the silencing effect of *skn-1* RNAi would be expected to depend primarily on RISC in the cytoplasm (37).

NRDE-4 is required for the recruitment of NRDE-1 to the targeted loci and subsequent deposition of the repressive H3K9me3 mark, which results in gene silencing. Furthermore, H3K9me3 has been implicated in transgenerational silencing of transgenes or endogenous loci mediated by exogenous RNAi (see ref. 18). These observations, and our findings that knockdown of *nrde-4* abolishes POE, led us to hypothesize that H3K9 methylation might function as a mediator of POE. Indeed, we found that treating F4 animals that showed POE with RNAi against *met-2* or *set-3*2, in contrast to their control siblings, eliminated POE in the F5 embryos (Fig. 4B). Although loss of MET-2 has been shown to enhance RNAi sensitivity (38), we found that neither *met-2* RNAi nor *set-32* RNAi significantly modifies the *skn-1(RNAi)* phenotypes of MY16 and JU1172 wild isotypes (p > 0.05; Fig. 4C). Thus, the loss of POE in the F5 generation with *met-2* or *set-32* RNAi is not attributable to modified RNAi response. We further found that a *set-32(−)* chromosomal mutation also eliminated TEI in N2/JU1491 crosses at the F2 generation, which implicates SET-32 not only in maintenance, but in initiation of this process. These results will be reported in greater depth elsewhere.

Importantly, we found that POE persisted in F5 embryos when a control gene, *ced-4*, was knocked down in the F3 and F4 generations. CED-4/Apaf-1 is involved in germline programmed cell death and has no known role in endoderm development or TEI (39). This result suggests that elimination of POE is not the result of a non-specific RNAi response (Fig. 4B and C, SI Appendix, Fig. S6). Collectively, these results suggest that, in response to dauer diapause, piRNAs in the germline direct histone methylation through the nuclear RNAi pathway, thereby maintaining POE across generations (Fig. 4D).

## Discussion

While massive epigenetic reprogramming ensures totipotency of the germline during animal development, some epigenetic marks escape erasure, leading to stable epigenetic inheritance that can persist through many generations. Such long-term epigenetic inheritance has the potential to provide a source of cryptic variation upon which evolutionary processes might act; however, little is known about natural epigenetic variation within a species, how it is influenced by environmental conditions, and the degree to which it influences GRN plasticity. In this study, we report four major findings that reveal cryptic natural epigenetic variation and its mechanistic action in modulating an embryonic GRN input in *C. elegans*: 1) dauer diapause can trigger POE that alters the SKN-1 requirement in endoderm development. 2) This effect is transmitted through the maternal germline across multiple generations apparently through nuclear signals. 3) Different combinations of wild isotypes exhibit variation in their capacity for establishing and maintaining these transgenerational epigenetic states. 4) This POE requires components of the piRNA-nuclear RNAi pathway and H3K9 trimethylation and does not appear to be eliminated by general RNAi response. These findings indicate that maintenance of an acquired epigenetic state in response to environmental stimulus can confer substantial plasticity to a core developmental program and may provide additional natural variation that may be subject to evolutionary selection.

### Dauer diapause induces persistent epigenetic inheritance

The perduring epigenetic effect that we have observed is triggered in parents that have experienced dauer diapause. Dauer entry and formation require extensive epigenetic remodeling and some of these changes are retained throughout the remainder of development: post-dauer adults contain distinct chromatin architecture and particular pools of small RNA that differ from animals that have not experienced dauer diapause (40). In addition, the progeny of starved animals show increased starvation resistance and lifespan (41, 42). Consistent with the model that the effect of ancestral developmental history is carried across generations, we found that dauer diapause leads to TEI that modifies the quantitative SKN-1 input in endoderm development. While it is conceivable that wild isotypes may exhibit natural variation in RNAi sensitivity and such difference can be epigenetically transmitted via maternal nuclei, we have previously shown that *skn-1* RNAi is highly effective across the wild isotypes (21), and we found fully penetrant *skn-1(RNAi)*-induced lethality in all experiments. In addition, all the strains we used in this study have been shown to be RNAi-competent (21, 43). It is therefore likely that the POE we report here reflects variation in endoderm specification *per se*, although it is conceivable that epigenetic influences on RNAi sensitivity may play a minor role.

The TEI we have observed varies in its long-term perdurance, depending on the wild isotypes involved. In crosses between the laboratory strain N2 and wild isotype JU1491, dauer diapause-induced POE lasted for three generations, but was subsequently lost, similar to the transmission dynamics of the silencing effect induced by exogenous RNAi (44). This progressive transgenerational loss in the effect may result from passive dilution of regulatory small RNAs and active restoration of an epigenetic “ground state” over generations, although the detailed mechanisms for such a process are not well understood (44). In contrast, in crosses between the MY16 and JU1172 wild isotypes, we observed stable TEI that lasted for at least 10 generations and which conceivably persists longer. Consistent with a recent study that identified genetic determinants of efficient germline maintenance and epigenetic reprogramming among *C. elegans* wild isotypes (9), our results showed that the generational duration of epigenetic inheritance may also be influenced by genetic background, suggesting an interplay between genetics and epigenetics (see also <u>MO</u>dified <u>T</u>ransgenerational <u>E</u>pigenetic <u>K</u>inetics (MOTEK) genes in ref. 45)

The transmission of the silencing effects of exogenous RNAi in *C. elegans* has provided an excellent paradigm for revealing mechanisms of epigenetic inheritance. In those studies, animals were subjected to RNAi targeting a transgene (e.g. *gfp*) or endogenous gene (e.g., *oma-1*) and the heritable RNAi response monitored over generations (e.g., ref. 8, 38, 46). While inheritance of exogenous RNAi and physiological responses triggered by changing environment share overlapping machinery, our results reveal two key differences between the two processes: 1) we demonstrated that epigenetic memory triggered by dauer diapause is transmitted exclusively through the maternal germline. This contrasts with the inheritance of exogenous RNAi (44) and transgenes (47), which show paternal bias. 2) We found that PRG-1 is required for the *maintenance* of POE. In contrast, the piwi-piRNA pathway had previously been shown to be required for the *initiation*, but not *maintenance* of transgene silencing in the germline. Once established, this silencing state depends on the nuclear RNAi pathway, which promotes deposition of H3K9me3 marks on the transgene (32, 33). Supporting our findings, however, Simon *et al.* demonstrated that PRG-1 is important for maintaining germline mortality through a mechanism that is independent from its action in transgene silencing. Animals that lack PRG-1 exhibit dysregulation of gene expression and reactivation of transposons and tandem repeats, showing that piRNAs are required to maintain silencing of at least some endogenous loci (48, 49).

### Relationship of POE to genomic imprinting in *C. elegans*

Genomic imprinting is perhaps the best-studied example of epigenetic inheritance. Differential DNA methylation or histone modifications on the two parental chromosomes, established during gametogenesis or post-fertilization, escape epigenetic reprogramming, causing genes to be expressed in a parent-of-origin manner (50, 51). However, in the case of *C. elegans*, animals that inherit the entire paternal genome are fertile and viable, as we and others have shown (26, 27). This observation reveals that genomic imprinting is not essential for normal development or survival in *C. elegans*, consistent with an early study in which animals containing individual chromosomes from only one parent were analyzed (52). Nevertheless, the X chromosome of sperm, unlike that of oocytes, is devoid of H3K4me2 activation marks, a pattern that persists through several rounds of cell division cycles during early embryogenesis (53). In addition, the expression of sperm-derived autosomal transgenes is greater than that in oocytes, which may result from differential epigenetic remodeling in sperm and oocyte chromatin upon fertilization (47). While these findings demonstrate the imprinting capacity of *C. elegans*, endogenous imprinted genes have not yet been reported.

We propose that passage through dauer diapause may induce paternal-specific silencing through deposition of repressive H3K9me3 on paternal loci that affect the SKN-1 requirement (21), leading to the POE we observed (Fig. 4D). Although imprinting is not essential for viability in *C. elegans*, its effects may become significant in response to environmental stimuli. For example, maternal dietary restriction elevates vitellogenin oocyte provisioning (54). Both yolk-associated fatty acids and small RNAs, which have been proposed to be associated with yolk particles, promote epigenetic changes in the nucleus, and might thereby direct establishment of parent-of-origin epigenetic marks (55). Dauer-favoring conditions also reduce insulin/insulin-like growth factor signaling and enhance starvation stress resistance in the progeny (54). With recent advances in techniques for examining transcriptional regulatory landscapes (e.g., ref. 51), it will be of interest to identify loci that are responsive to environmental stimuli and that may be differentially imprinted across generations.

### The potential role of cryptic epigenetic variation in accelerating evolutionary change

It is clear that in *C. elegans*, stress responses can be transmitted transgenerationally and influence physiology adaptively in the offspring (e.g. refs. 41, 42). In *Arabidopsis*, it has been shown that experimentally induced, or naturally occurring epigenetic variations, once stabilized, can be subjected to artificial selection (56, 57), highlighting the potential capacity of TEI to facilitate adaptation and evolution. While TEI is prominent in worms and plants with a short life cycle, and hence environmental conditions may be relatively constant through multiple generations, there is evidence that TEI may also occur in mammals with much longer reproductive cycles (58). Epigenetic inheritance may be especially important in organisms with low genetic diversity, such as those, including *C. elegans* and *Arabidopsis*, that propagate by self-fertilization. Many *C. elegans* strains isolated from neighboring locations are near-identical and polymorphism rates are low even among genetically distinct isotypes (59). In such homogenous, genetically non-diverse populations, epigenetic variations may provide a particularly rich resource upon which natural selection may act.

Environmental factors can induce plastic phenotypic changes that are subjected to Darwinian selection. Over time, the phenotypic variants may become genetically fixed, a process known as “genetic assimilation” (23). As the rate of genetic mutations is low in *C. elegans* (2.1 × 10^−8^ per nucleotide site per generation) (60), heritable epigenetic variants may act as a buffer to cope with rapid environmental change before adaptive mutations arise. Alterations in epigenetic states can also affect mutation rates and trait evolution (61) and TEI might therefore accelerate the rate of evolution by facilitating genetic assimilation. We propose that epigenetic inheritance affecting SKN-1 dependence may contribute toward the rapid change in the modulation of the endoderm GRN that we previously observed among *C. elegans* wild isotypes (21). We note that that the POE we observed likely reflects changes in epigenetic states that occur in response to a stressful environment (starvation and dauer formation) even in the absence of mating between wild isotypes (41). That is, we propose that crossing is not necessary for implementation of the effect; rather, crosses between strains with distinct *skn-1(−)* phenotypes primarily provides a readout that allows the underlying TEI to be revealed. SKN-1 acts pleiotropically, including its roles in mesendoderm specification, oxidative stress and unfolded protein responses, promoting longevity, and modulating metabolism during starvation (reviewed in ref. 62). It is conceivable that SKN-1 is particularly susceptible to plastic changes in its regulatory outputs as a means of adapting to frequently varying environmental conditions. It is possible that pleiotropically acting genes involved in stress response pathways, perhaps those that also modulate dauer development, may play a cryptic role in endoderm development. Hence, (heritable) change of function or expression of those genes in response to stress may subsequently alter the signaling during endoderm specification. In the wild, *C. elegans* experiences a boom-and-bust cycle and most worms isolated in the wild are present in the dauer stage (63). As we have shown, dauer diapause is associated with strong heritable epigenetic effects (41) that may, therefore, influence developmental plasticity and adaptive evolution in response to the local environment. We believe that our findings may provide among the first example of environmentally-induced heritable epigenetic changes that modulate developmental input(s) into an embryonic GRN.

## Materials and Methods

Refer to SI Appendix for full methods.

### *C. elegans* strains

N2 (Bristol, UK), MY16 (Mecklenbeck, Germany), JU1491 (Le Blanc, France), and JU1172 (Concepcion, Chile). JR3336*, (elt-2∷GFP) X; (ifb-2∷GFP) IV*. PD2227 (27), *oxIs322 II; ccTi1594 III. oxIs322 contains [myo-2p∷mCherry∷H2B + myo-3p∷mCherry∷H2B + Cbr-unc-119(+)] II. ccTi1594* contains *[mex-5p∷GFP∷gpr-1∷smu-1 3’UTR + Cbr-unc-119(+), III: 680195] III.* MD701, *bcIs39 [lim-7p∷ced-1∷GFP + lin-15(+)] V*. JR3666, PD2227 and MD701 are N2-derived transgenic animals.

### Dauer induction and POE assays

The animals were maintained on NGM plates seeded with OP50. Once the cultures became crowded and exhausted their food supply, they were incubated for an extra two weeks at 20°C. The worms were then washed with M9 buffer and incubated in 1% sodium dodecyl sulfate (SDS) for 30-60 minutes with gentle agitation to select for dauer larvae (64). Isolated dauer larvae were then washed with M9 to remove all SDS and allowed to recover overnight on 60 mm NGM plates seeded with OP50.

Reciprocal crosses were set up using L4s and the animals were allowed to mate overnight. Control experiments using well-fed animals were performed in parallel. Mated hermaphrodites, as indicated by the presence of copulatory plugs (except for crosses involved N2 males which do not deposit plugs), were transferred to a fresh NGM plate to lay eggs for ~5-7 hours and the resulting early brood was discarded to avoid contamination of self-progeny. The hermaphrodites were then transferred to a fresh seeded NGM plate to lay eggs overnight. The hermaphrodite (P0) was then removed, leaving the F1s alone. Once the F1 worms reached early or mid-adulthood, they were treated with 15% alkaline hypochlorite solution to obtain F2 embryos which were allowed to hatch on food. This procedure was repeated until the F4 generation (F10 for the experiment shown in Fig. 1D) was obtained. At each generation, L4 worms were used to analyze *skn-1* RNAi phenotype. As SKN-1 is maternally provided, the F2 phenotype, for example, is the result of *skn-1* RNAi treatment of F1 mothers. To examine the maternal effect, mating was performed on *skn-1* RNAi plates. The arrested F1 embryos from mated hermaphrodites were then collected for quantification of the number of animals containing gut granules (Fig. 1A).

For crosses between PD2227 hermaphrodites and JU1491 males, the POE assay was performed as described above. F1 L4s were immobilized on a 5% agar pad with 5 mM levamisole diluted in M9 and observed using Nikon Eclipse Ti-E inverted microscope. Mosaic worms that expressed *myo-2*∷mCherry, but not *myo-3*∷mCherry and *mex-5*∷GFP (i.e., PD2227-derived AB and JU1491-derived P_1_), were recovered on a seeded NGM plate in the presence of M9. 20 F2 animals were then randomly selected and observed to ensure no worms expressed fluorescent markers, i.e. JU1491 nuclear genotype (Fig. 2D, SI Appendix, Fig. S6).

### RNAi

Feeding-based RNAi experiments were performed as described (21). RNAi clones were obtained from either the Vidal (65) or Ahringer (66) libraries. RNAi bacterial strains were grown at 37°C in LB containing 50 μg/ml ampicillin. The overnight culture was then diluted 1:10. After four hours of incubation at 37°C, 1 mM IPTG was added and 60-100 μl was seeded onto 35 mm agar plates containing 1 mM IPTG. Seeded plates were allowed to dry and used within five days when kept at room temperature. For *skn-1* RNAi, five to 10 L4 animals were placed on each RNAi plate. 24 hours later, they were transferred to another RNAi plate to lay eggs for 12 hours. The adults were then removed, leaving the embryos to develop for an extra 5-7 hours. Embryos expressing birefringent gut granules were quantified and imaged on an agar pad using a Nikon Ti-E inverted microscope under dark field with polarized light (SI Appendix, Fig. S1B). We note that the *skn-1* sequence is highly conserved in the isotypes used in this study. It is therefore highly unlikely that variation in *skn-1(RNAi)* phenotypes is attributable to underlying *skn-1* polymorphisms.

For *met-2*, *set-32*, *nrde-4*, *prg-1/2* and *ced-4* RNAi, 15-30 F3 L4 animals showing POE were placed on plates of *E. coli* containing an empty control vector (L4440) or expressing double-stranded RNA. 24 hours later, they were transferred to another RNAi plate and allowed to lay eggs for another ~7 hours. The adults were then removed and the F4 animals were allowed to develop on RNAi feeder bacteria. F4 L4 larvae were used for the *skn-1* RNAi assay for POE as described above (Fig. 4A).

## Acknowledgments

We apologize to colleagues whose work could not be cited due to limited space. We thank Coco Al-Alami for experimental assistance at early stages of the project. We thank members of the Rothman lab, especially Dr. Pradeep M. Joshi, for helpful advice and feedback. Nematode strains used in this work were provided by the Caenorhabditis Genetics Center, which is funded by the National Institutes of Health − Office of Research Infrastructure Programs (P40 OD010440). This work was supported by grants from NIH (#1R01HD082347 and # 1R01HD081266). YNTC is supported by the Norwegian Research Council (#250049).

## Supplementary Materials and Methods

### Worm culture

Worm strains were maintained as described (1) and all experiments were performed at 20°C unless noted otherwise. To ensure no carry-over of a parental stress response, a fresh worm stock was obtained from −80°C and maintained on 150 mm NGM plates seeded with *E. coli* OP50 for at least five generations prior to beginning experiments. To avoid genetic drift and lab domestication, a fresh worm stock was obtained every ~30 generations.

To obtain males for crosses, 20–30 L4 hermaphrodites were picked into 7% ethanol solution in microcentrifuge tubes and rotated for one hour (2). Worms were pelleted by centrifugation at 2000 rpm for 30 seconds. They were then transferred to NGM plates seeded with *E. coli* OP50. F1 male progeny were mated with sibling hermaphrodites to establish male stocks.

### Viability and embryonic lethality scoring

To score viability, young hermaphrodites (F1 progeny of the reciprocal crosses) were allowed to lay eggs on an NGM plate seeded with OP50. The next day, newly hatched L1s (F2) were transferred to individual seeded plates. “Larval lethal” was defined as the percentage of worms that arrested as L1s. Worms that reached adulthood but failed to reproduce in five days were scored as sterile. To score embryonic lethality, individual young hermaphrodites (F1) were allowed to lay eggs on an NGM plate seeded with a small drop of OP50 for ~4-8 hours. The hermaphrodites were then removed, leaving the embryos. The fraction of unhatched embryos were counted and scored ~24 hours later. At least two independent broods were scored. We note that there were no obvious differences in brood sizes between the various crosses.

### Statistics and figure preparation

Statistics were performed using R software v3.4.1 (https://www.r-project.org/). Two-sample two-tailed t-tests were used to compare the *skn-1* RNAi phenotype between two groups, unless stated otherwise. Welch’s t-tests were performed if the variances of the two groups being compared were not equal. Fisher’s exact tests were used to determine the difference in proportions (Fig. 3B, C; Fig. 4C). Plots were generated using R package ggplot2. Figures were assembled in Inkscape v0.92.4 (https://inkscape.org/).

## Supplementary Figures and Table

**Fig. S1.**
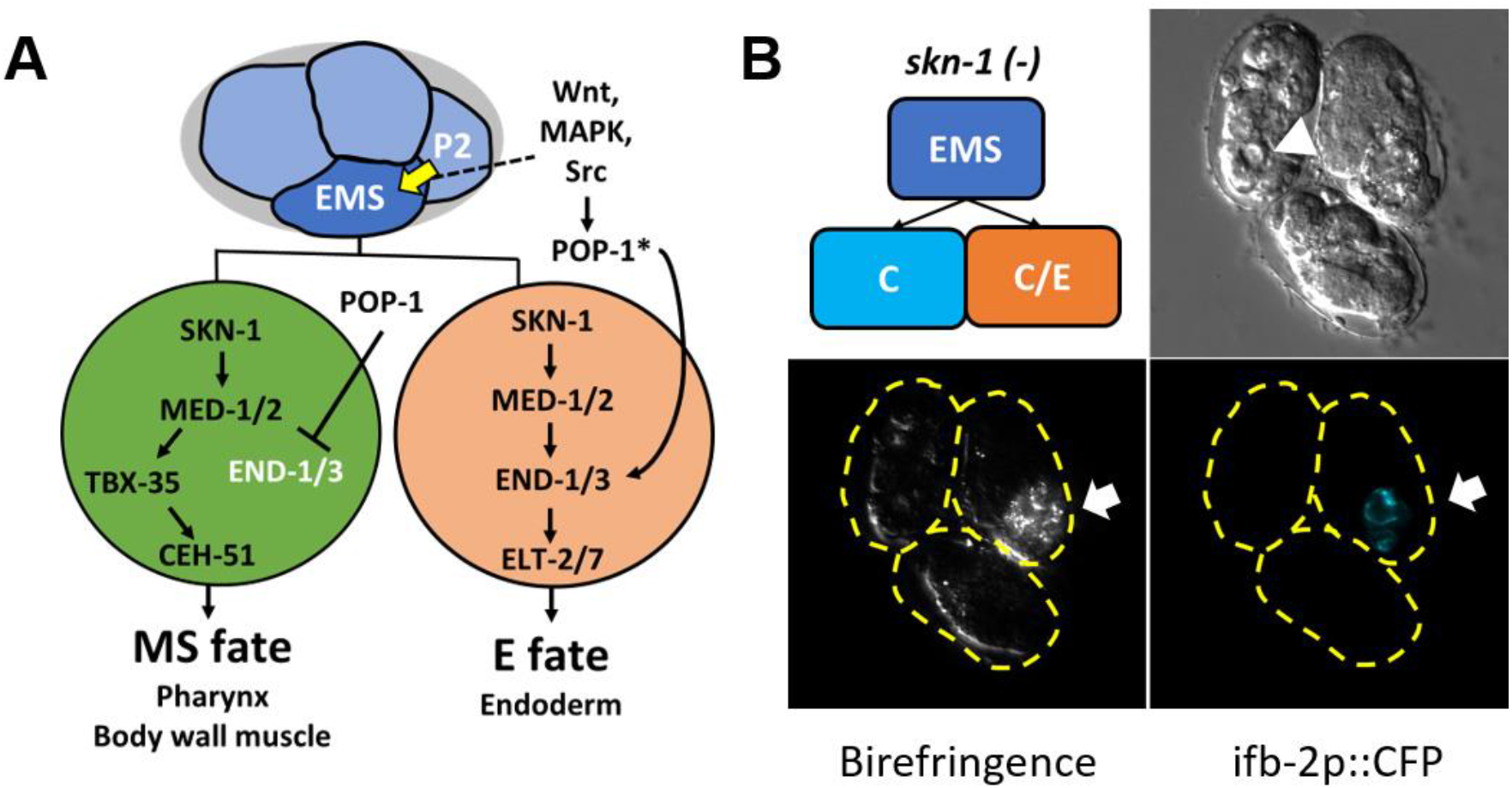
Endoderm development in *C. elegans.* A) In the four-cell stage, signaling from the posterior P_2_ cell (Wnt, MAPK and Src) polarizes the blastomere, leading to POP-1 asymmetry in the descendants of EMS with high levels of nuclear POP-1 in anterior MS and low levels of nuclear POP-1 in the posterior E cell. In the anterior MS cell, high nuclear POP-1 represses *end-1/3*, allowing SKN-1 to induce the MS fate though the actions of TBX-35 and CEH-51 (3, 4). In the posterior E cell, which remains in contact with P_2_, POP-1 is converted to an endoderm activator (POP-1*) and along with SKN-1 activates *end-1/3* and *elt-2/7* genes, resulting in the endoderm fate. B) Loss of *skn-1*, either by RNAi or in loss-of-function mutation, results in 100% of the embryos to arrest as MS is mis-specified to form a C blastomere. In some embryos, E also converts to C. The arrowhead indicates internal cuticle-lined cavity in *skn-1(−)* embryos, resulting from the E to C transformation. The E fate is retained in some embryos as indicated by the presence of gut-specific *ifb-2*∷CFP expression and birefringent gut granules (arrows), owing to the partially redundant input of POP-1*.

**Fig. S2.**
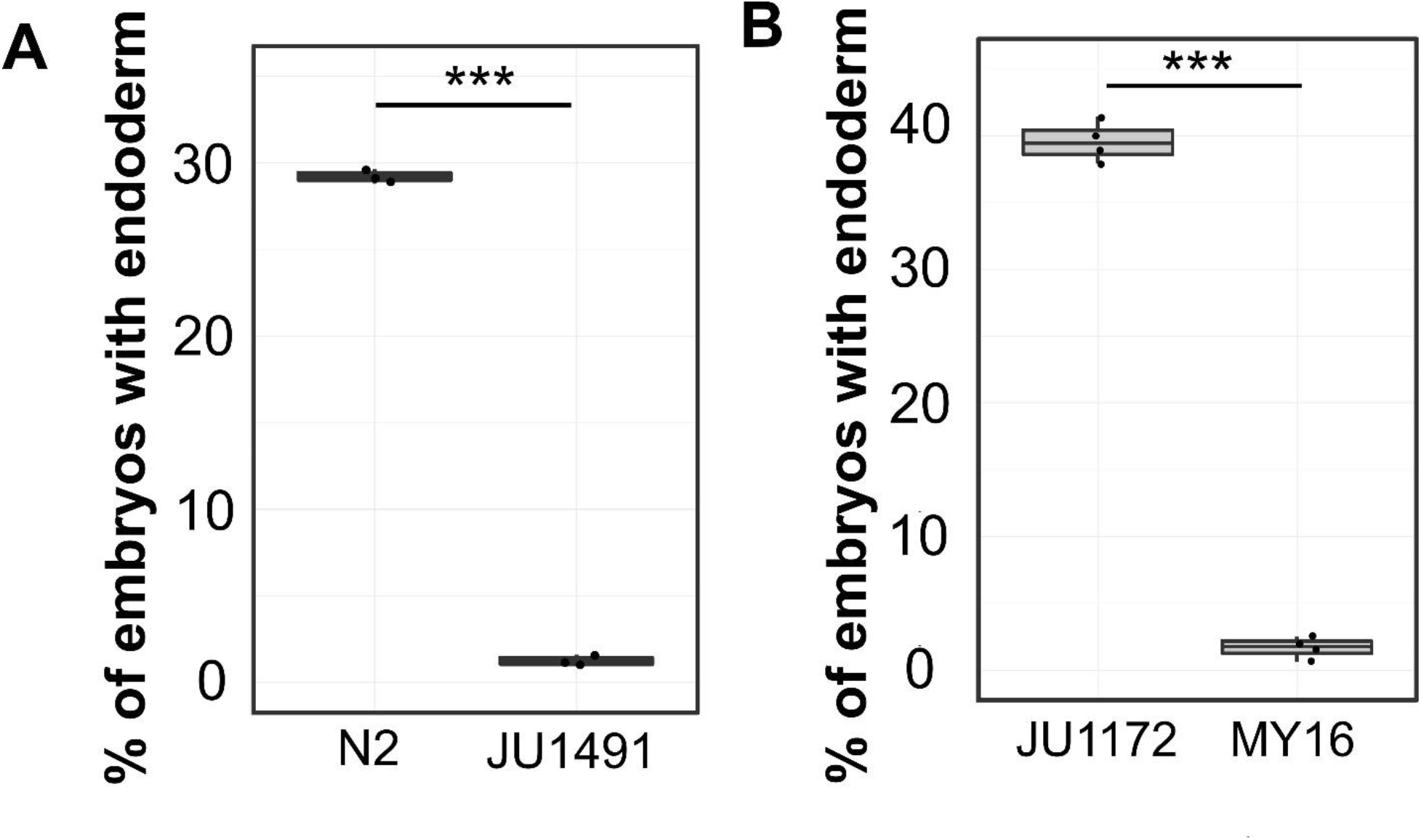
Natural variation in SKN-1 requirement. A) *skn-1(RNAi)* phenotype of laboratory strain N2 and wild isotype JU1491. B*) skn-1(RNAi)* phenotype of wild isotypes JU1172 and MY16. At least three replicates were performed for each strain, with >200 embryos per replicate. Two sample t-test (*** p-value ≤ 0.001).

**Fig. S3:**
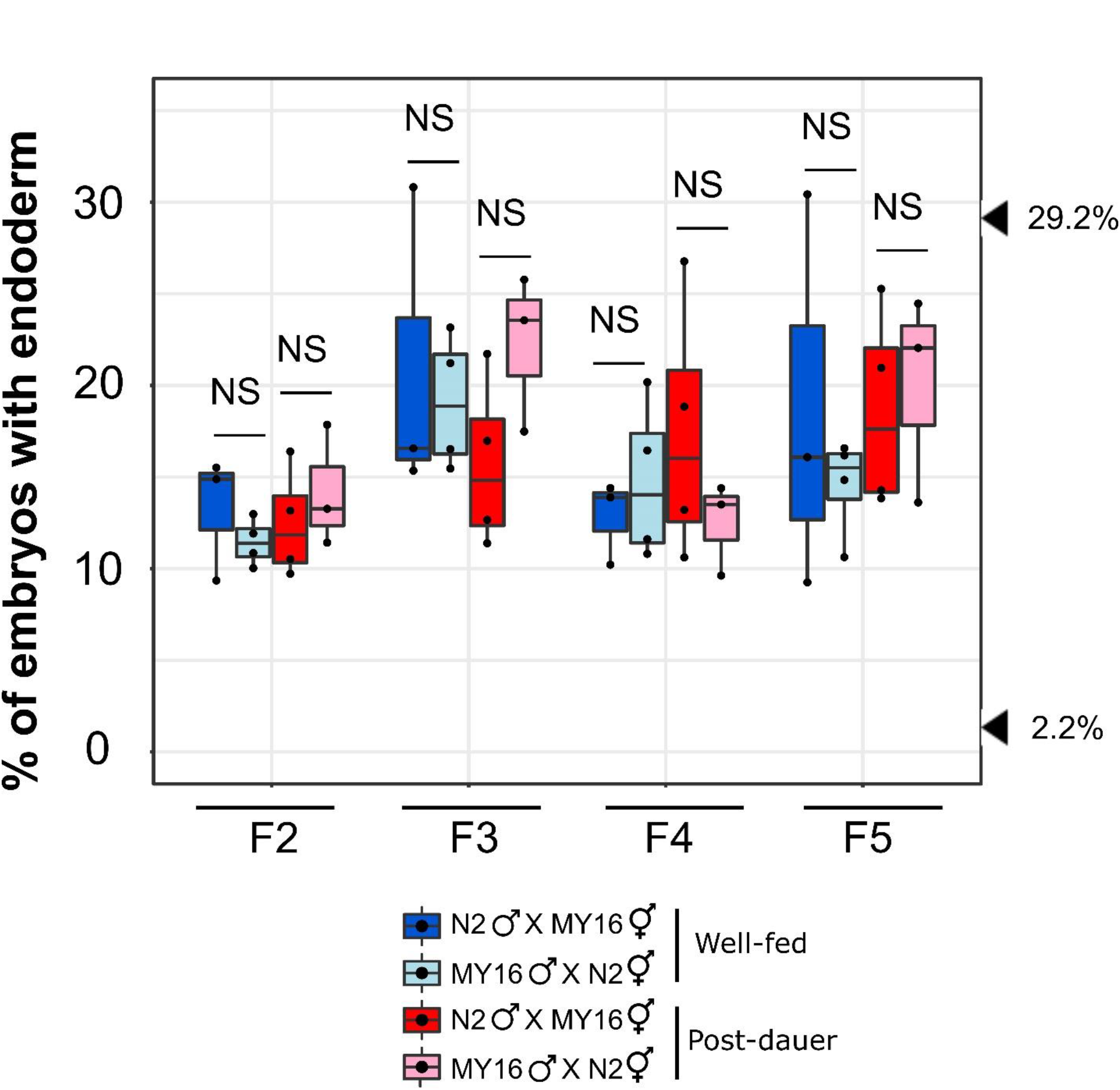
Absence of POE in N2 × MY16. No POE was observed in *skn-1(RNAi)* embryos from four generations: F2, F3, F4 and F5 derived from N2 × MY16 crosses. At least three independent crosses were performed for each treatment. Arrowheads indicate the phenotypes of JU1172 (40.0%) and MY16 (2.2%). Two sample t-test (NS p > 0.05). Boxplot represents median with range bars showing upper and lower quartiles.

**Fig. S4:**
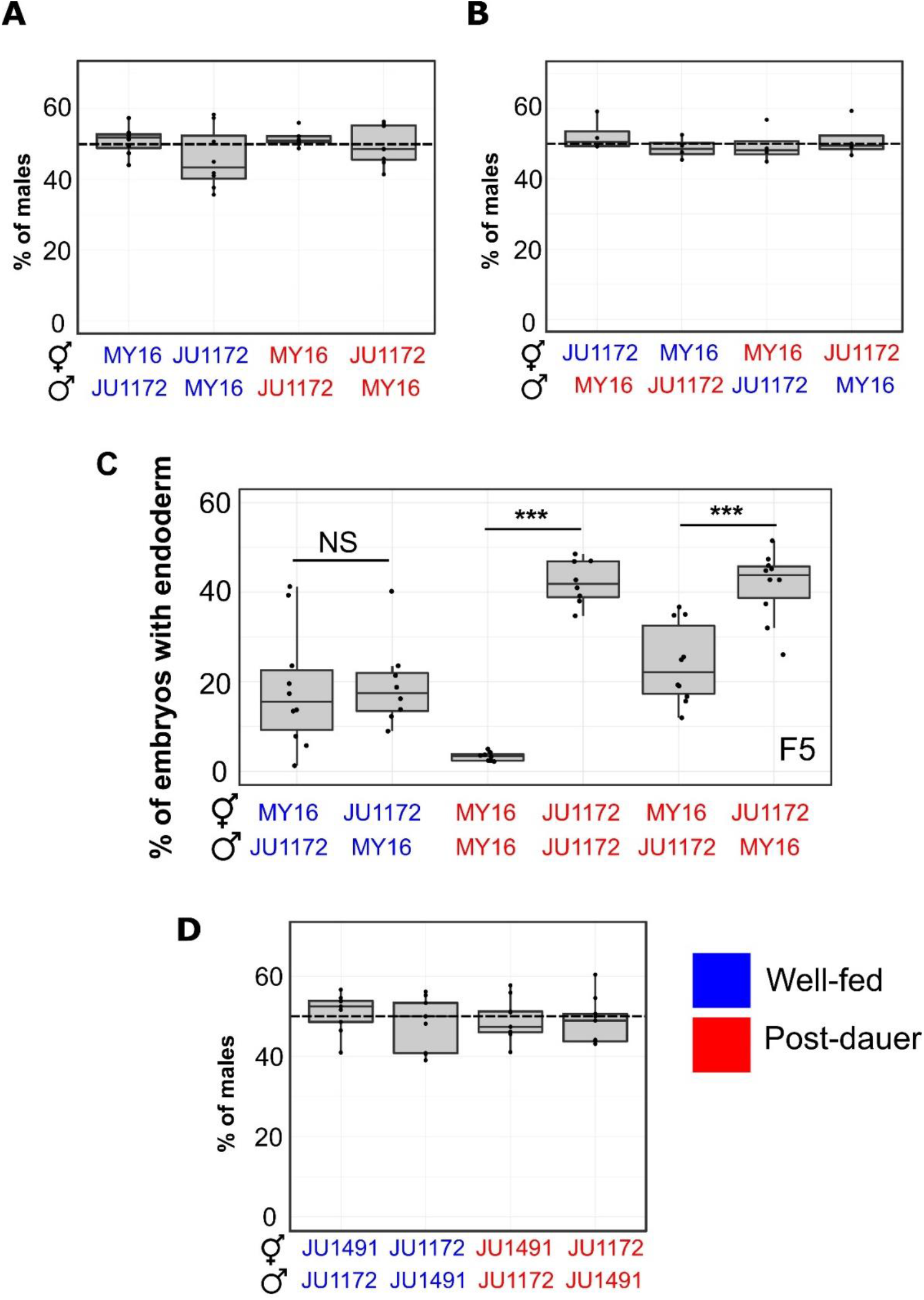
POE is not the result of frequent selfing. A and B) Male frequency in the F1 offspring from reciprocal crosses between MY16 × JU1172 (related to Fig. 1D and E). In all cases, ~50% of males were found (one-sample t-test, p > 0.05). C) Individual data points represent *skn-1(RNAi)* F5 embryos of lines derived from isolated F1s. Two sample t-test (NS p-value > 0.05, *** p-value ≤ 0.001). D) Male frequency in the F1 offspring from reciprocal crosses between JU1172 × JU1491 (related to Fig. 2C). Boxplot represents median with range bars showing upper and lower quartiles. Blue text represents experiments performed using well-fed animals while red text represents experiments performed using post-dauer animals.

**Fig. S5:**
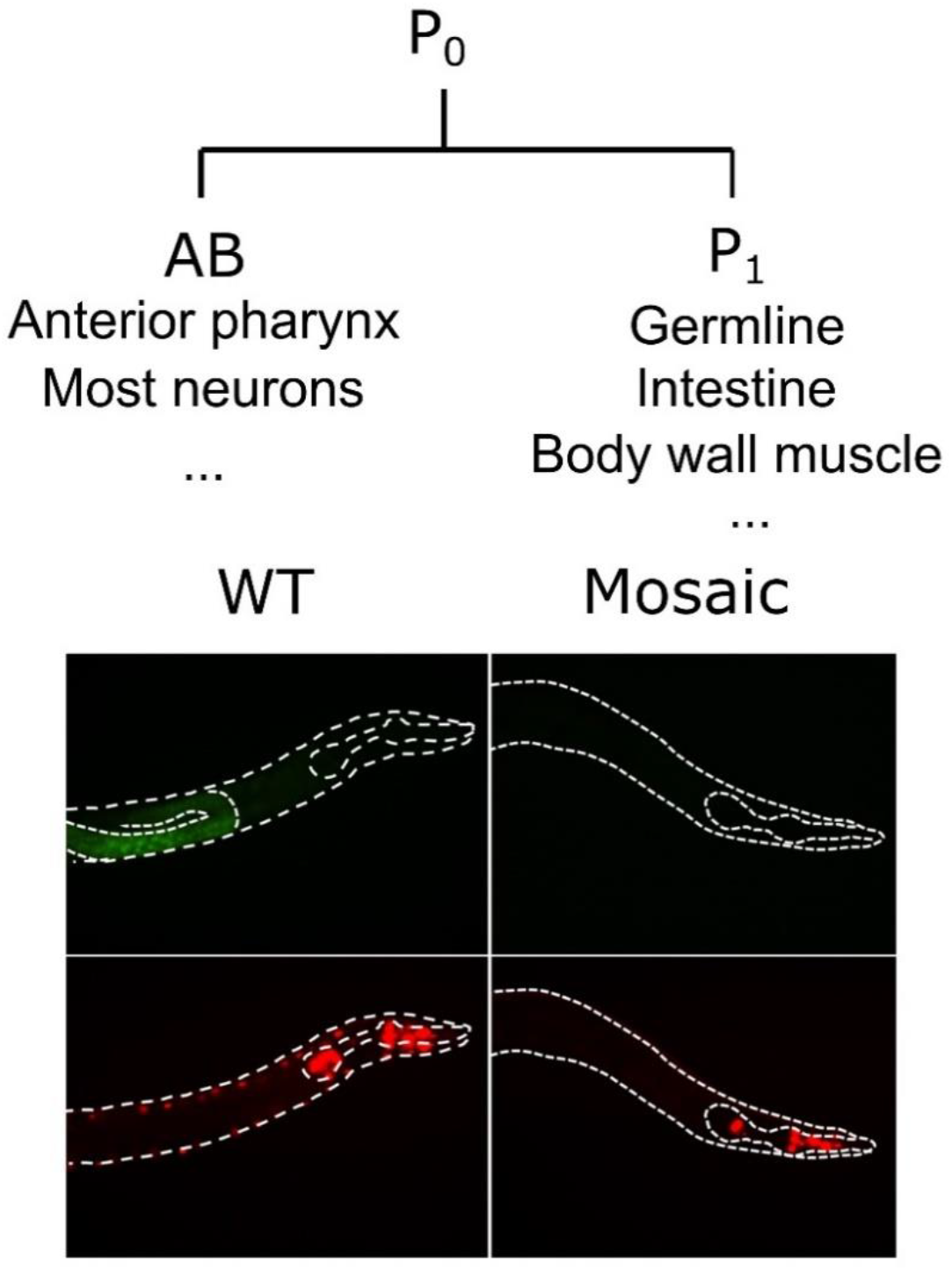
Non-Mendelian chromosomal segregation in GPR-1 overexpression animals. The two daughter cells, AB and P_1_, arising from the first asymmetrical cell division develop into different cell lineages. When PD2227 (N2^GPR-1 OE^) hermaphrodites are mated to wild-type JU1491, unequal segregation of maternal and paternal chromosomes occurs, causing AB to contain only the maternal chromosomes and P_1_ to contain only the paternal chromosomes. F1 mosaic animals express the maternally-derived pharyngeal marker (*myo-2*∷mCherry) but not the body wall muscle (*myo-3*∷mCherry) and germline (*mex-5*∷GFP) markers (5).

**Fig. S6:**
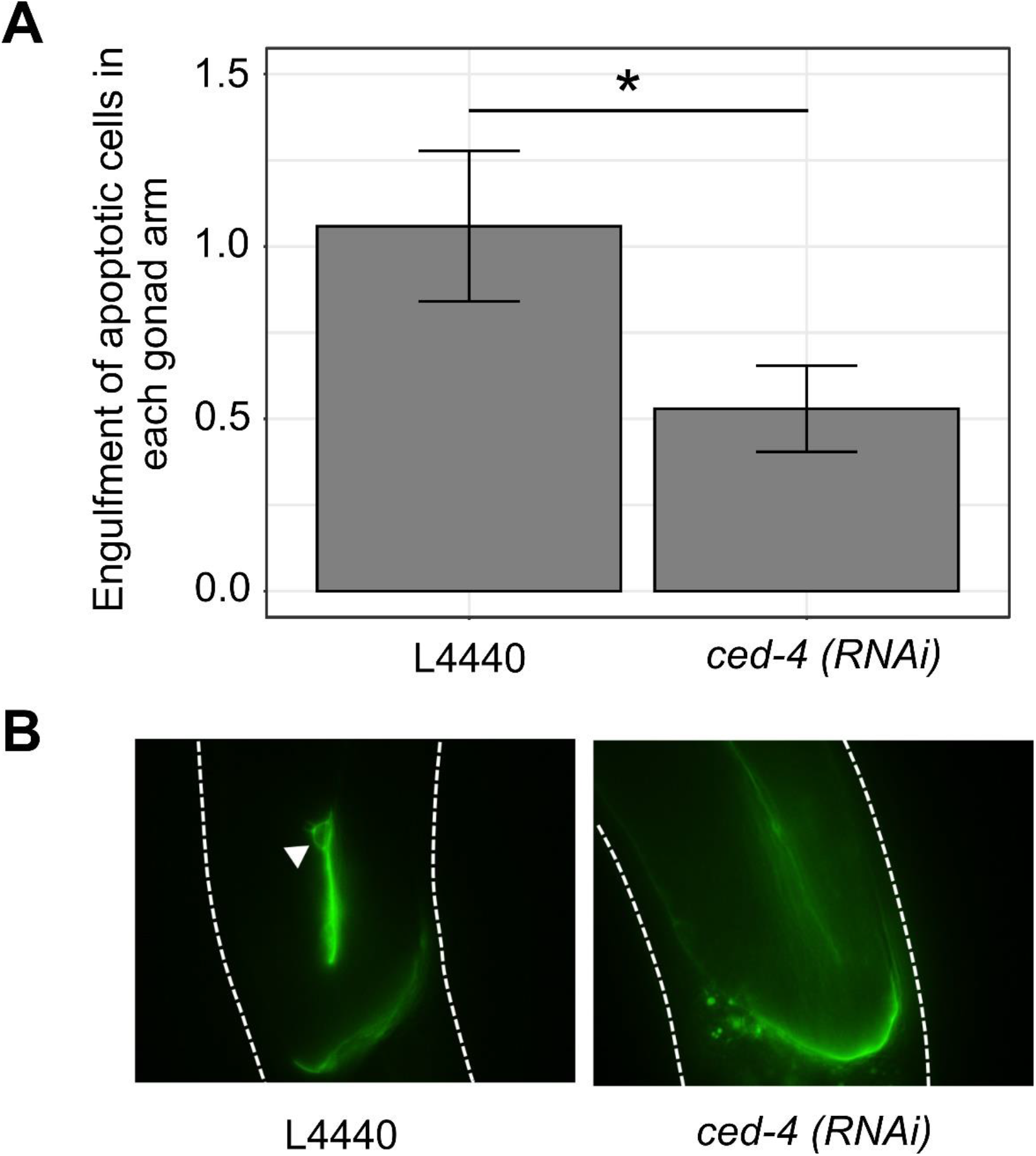
*ced-4* RNAi suppresses apoptotic cell death of meiotic germ cells. A) Quantification of *bcls39* engulfment of apoptotic cell corpses in MD701, which is an N2-derived strain. L4 animals were placed on plates of *E. coli* containing an empty control vector (L4440) or expressing double-stranded RNA. 24 hours later, they were transferred to another RNAi plate and allowed to lay eggs for ~7 hours. The adults were then removed and the F1 animals were allowed to develop on RNAi feeder bacteria. The number of apoptotic germ cells were scored in young adults (n=17 for each treatment). Error bars represent +/− standard error of the mean. Two sample t-test (* p-value ≤ 0.05). B) Representative images of control and *ced-4(RNAi)* worms. *bcls39* [CED-1∷GFP] clusters around apoptotic cell corpses are indicated by white arrowhead.

**Table S1.** Polymorphisms in the mitochondrial protein coding genes between MY16 and JU1172 (6).

| **Genes** | **Position** | **MY16** | **JU1172** | **Modifications** |
| --- | --- | --- | --- | --- |
| ***nduo-6*** | 112 | T | G | Val4Leu |
|  | 227 | G | A | Val39Ile |
|  | 382 | A | T | Leu90Phe |
| ***nduo-1*** | 2,476 | A | T | Lys238Asn |
| ***atp-6*** | 3,230 | GTAATA | GA | Frameshift |
| ***nduo-2*** | 3,614 | G | T | Cys66Phe |
|  | 3,997 | G | A | Gly194Ser |
| ***ctb-1*** | 5,365 | A | G | Ile288Val |
|  | 5,367 | T | A | Ile288Met |
| ***ctc-1*** | 9,242 | T | A | Ile466Met |
| ***ctc-2*** | 10,003 | T | C | Leu119Met |
| ***nduo-5*** | 12,714 | G | A | Ala342Thr |
|  | 12,906 | G | A | Val406Met |

